# Transcriptomic Analysis of the Innate Immune Response to *in vitro* Transfection of Plasmid DNA

**DOI:** 10.1101/2021.06.21.449271

**Authors:** Eric Warga, Matthew Tucker, Emily Harris, Jacob Elmer

## Abstract

The innate immune response to cytosolic DNA is intended to protect the host from viral infections, but it can also inhibit the delivery and expression of therapeutic transgenes in gene and cell therapies. The goal of this work was to use mRNA-sequencing to reveal correlations between the transfection efficiencies of four cell types (PC-3, Jurkat, HEK-293T, and primary CD3^+^ T cells) and their innate immune responses to nonviral gene delivery. Overall, the highest transfection efficiency was observed in HEK-293T cells (87%), which upregulated only 142 genes with no known anti-viral functions. Lipofection upregulated a much larger number (n = 1,057) of cytokine-stimulated genes (CSGs) in PC-3 cells, which also exhibited a significantly lower transfection efficiency. However, the addition of serum during Lipofection and electroporation significantly increased transfection efficiencies and decreased the number of upregulated genes in PC-3 cells. Finally, while Lipofection of Jurkat and Primary T cells only upregulated a few genes, several anti-viral CSGs that were absent in HEK and upregulated in PC-3 cells were observed to be constitutively expressed in T cells, which may explain their relatively low Lipofection efficiencies (8-21%). Indeed, overexpression of one such CSG (IFI16) significantly decreased transfection efficiency in HEK cells to 33%.

## INTRODUCTION

In recent years, multiple gene therapy treatments using adeno-associated viruses (AAVs) and lentiviruses (LVs) have been shown to effectively treat a variety of diseases in the clinic.^1^ For example, AAV is used to deliver a functional copy of the RPE65 gene to patients with Leber’s Congenital Amaurosis (LCA) in Luxturna®,^2^ while Zolgensma® uses an AAV to deliver the SMN1 gene to patients with spinal muscular atrophy (SMA). Likewise, delivery of the chimeric antigen receptor (CAR) to T cells is done with an LV in three different CAR-T cell therapies for B-cell lymphomas (Kymriah®, Yescarta®, and Tecartus™).^3–5^ Several other promising gene therapies for hemophilia, Duchenne’s muscular dystrophy (DMD), immunodeficiency, and Pompe disease are also progressing through clinical trials.^6–9^ All of these therapies have demonstrated that gene delivery can be a lifesaving treatment for patients where other treatment mechanisms have failed.^4^ However, several issues have arisen in regards to LV and AAV vehicles, including high treatment costs (e.g., $2M for Zolgensma®), safety concerns, and relatively low transduction efficiencies in some patients.^2,4,10–16^

These issues with viral gene delivery vehicles have motivated a growing demand for safer and less expensive nonviral gene delivery methods. Several nonviral gene delivery vehicles and methods have been developed, including Lipofectamine, polyethyleneimine (PEI), lipid/polymer hybrids, nanoparticles, and electroporation.^1,17–19^ However, while these methods are generally less expensive and potentially safer than AAVs and LVs, they tend to provide lower transfection efficiencies than viruses.^20,21^ Consequently, improving the efficiency of nonviral transgene delivery and expression is an important and highly active area of research.

One possible way to accomplish this is by inhibiting the innate immune response to foreign genes, which is shown in Figure 1. For example, both viral and nonviral transgenes can be detected by endosomal or cytosolic DNA sensors (e.g., IFI16), which then utilize adaptor proteins like STING to trigger a signaling cascade of kinases and transcription factors that culminates in the activation of cytokines (e.g., interferon *λ*). These cytokines activate additional pathways that induce the expression of cytokine-stimulated genes (CSGs) like IFIT1 and OAS1 that can directly inhibit the delivery and expression of viral and nonviral transgenes.^22–24^ For example, IFI16 has been shown to decrease plasmid-driven transgene expression by directly binding and blocking viral promoters.^25,26^ In addition, IFI16 is also a cytosolic/nuclear dsDNA sensor that amplifies the innate immune response by activating STING and IRF3.^27–29^

**Figure 1.**
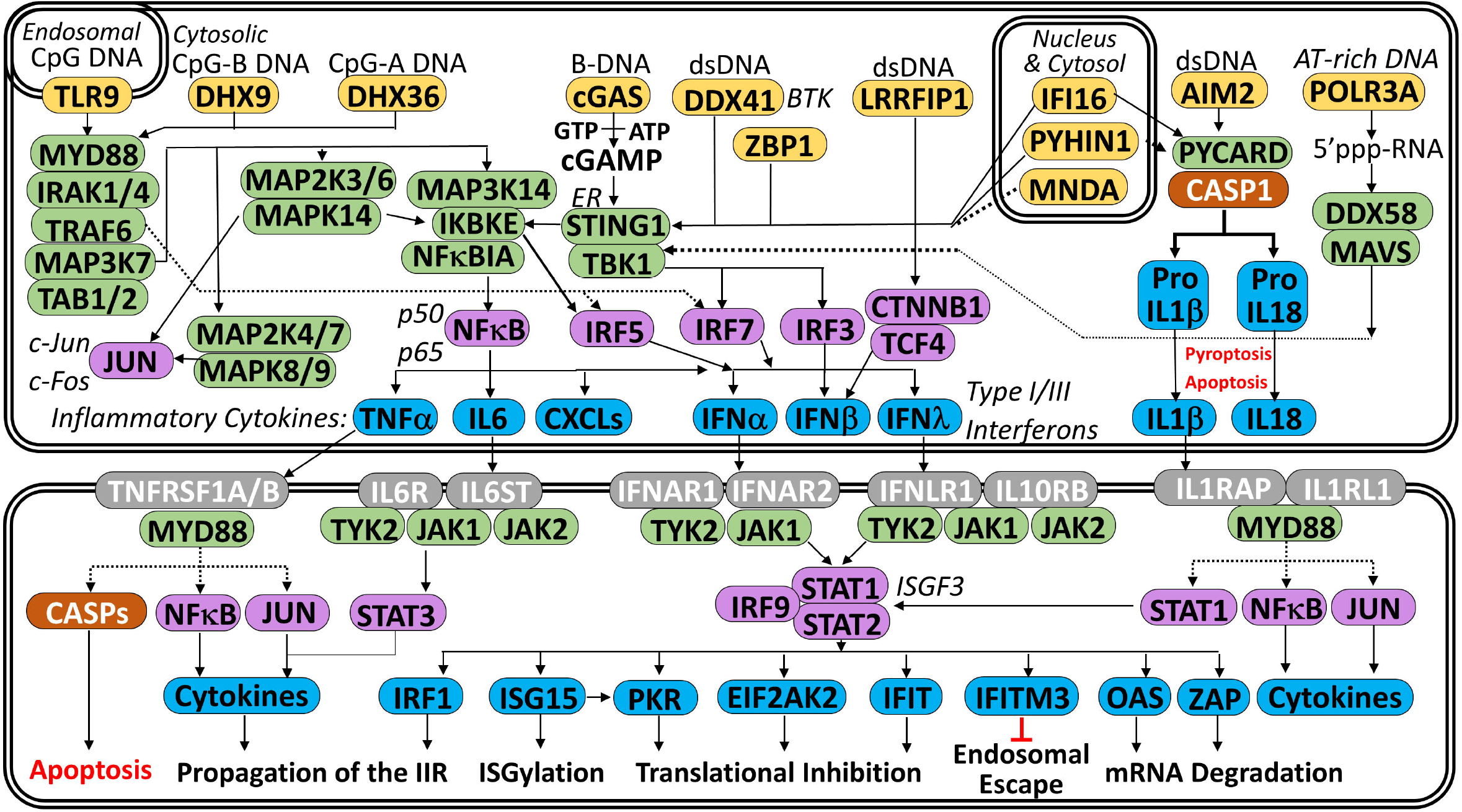
Plasmid DNA can be recognized by several DNA sensors (yellow), which trigger a cascade of kinases (green) and transcription factors (purple) that culminates in the expression of cytokines that are secreted and bind to cognate receptors (gray) that induce the expression of cytokine stimulated genes (blue) that can trigger apoptosis or inhibit transgene expression in a variety of ways.

While many studies have focused on specific components of the innate immune response to plasmid DNA and nonviral gene delivery, our overall knowledge of the transcriptomic profile of different cell types following transfection is still incomplete. In this study, mRNA-sequencing was used to elucidate the innate immune response to plasmid DNA at the transcriptome level in a panel of 4 cell lines (HEK293T, PC-3, Jurkats, and Primary T Cells) with varying transfection efficiencies to identify host cell genes that may inhibit transfection. In addition, the effect of serum and electroporation on the particularly potent innate immune response of PC-3 cells was also investigated.

## RESULTS

### Transfection Efficiency

Three cancer cell lines (HEK-293T, PC-3, and Jurkat T cells) with a range of different Lipofection efficiencies (see Figure 2A) were selected for this study with the goal of identifying correlations between their wide range of transfection efficiencies and gene expression patterns. Primary T cells were also included in the study because they are used for chimeric antigen receptor (CAR) T cell therapy. As shown in Figure 2A, Lipofection of pDNA into HEK-293T cells in serum-free media (SFM) provides a relatively high transfection efficiency (87.3 ± 1.2% GFP^+^ cells; measured at 24 hours post-transfection), while progressively lower transfection efficiencies were observed for PC-3 (46.3 ± 3.7% GFP^+^ cells) and Jurkat T cells (21.2 ± 3.4%). Finally, despite optimization of a Lipofection protocol in serum-free X-VIVO15 media, primary T cells exhibited the lowest Lipofection efficiency (8.1 ± 0.8%).

**Figure 2.**
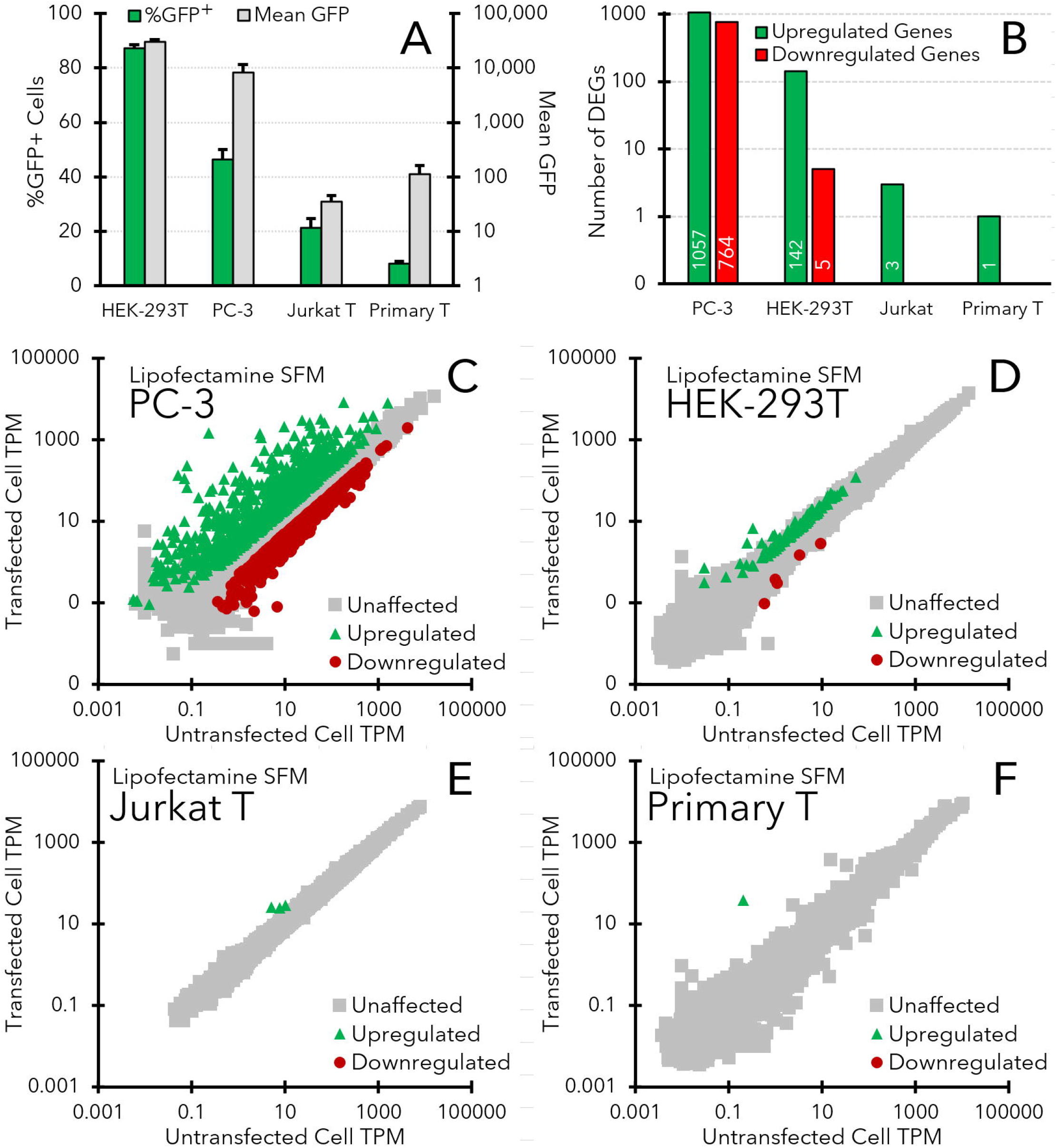
Differences in transfection efficiency and transcriptomes between cell lines transfected with Lipofectamine LTX in serum-free media (SFM). (A) Transfection efficiency (%GFP^+^ cells) and transgene expression levels (Mean GFP) for each cell line at 24 hours after transfection with Lipofectamine LTX and pEF-GFP plasmid DNA in serum free media (SFM). Representative flow cytometry histograms and fluorescent microscopy images for each cell line are also shown in Figure S1. (B) Total number of upregulated and downregulated differentially expressed genes (DEGs) observed in each cell line following Lipofection in SFM. DEGs were defined as having at least a 2-fold change in TPM that was statistically significant (P_adj_ < 0.05) over the course of 3 independent experiments. (C-F) Gene expression levels (TPMs) in HEK-293T cells (C), PC-3 cells (D), Jurkat T cells (E), and primary CD3^+^ T cells (F) that were either transfected with Lipofectamine LTX (y-axis) or not transfected (x-axis). All TPM values are averaged from 3 separate transfections and corresponding mRNA-sequencing experiments. Green triangles indicate genes that were significantly upregulated in transfected cells (P_adj_ < 0.05), red circles indicate downregulated genes (P_adj_ < 0.05), and gray squares represent unaffected genes.

A similar downward trend was also observed in GFP expression levels between the cell lines (Figure 2A). While GFP was expressed at high levels in HEK-293T cells (Mean GFP = 30,266 ± 2,715) that were brightly fluorescent (Figure S1C), GFP levels were significantly lower in PC-3 cells (Mean GFP = 8,332 ± 3,239) and extremely low in Jurkat and CD3^+^ Primary T cells (Mean GFP = 35 ± 11 & 112 ± 50, respectively). Fluorescent microscopy images shown in the supplementary information (Figure S1C-F) concur with these measurements, showing a decrease in the brightness and number of fluorescent cells between the HEK-293T cells and the other cell lines.

### Differential Gene Expression

A graphical summary of the number of genes that were differentially expressed between the control and transfected samples for each cell line is shown in Figure 2B, while complete lists of the DEGs are included in the supplementary information. Figure 2 also shows specific TPM plots for PC-3 (Figure 2C), HEK-293T (Figure 2D), Jurkat T cells (Figure 2E), and primary T cells (Figure 2F).

Overall, PC-3 cells exhibited the strongest innate immune response, in which 1,057 genes were upregulated and 764 genes were downregulated after Lipofection in SFM (Figures 2B/C). In contrast, the HEK-293T cells that exhibited a higher transfection efficiency than PC-3 cells upregulated a much lower number of genes (n = 142), with only 10 genes that were upregulated more than 4-fold (Figure 2D). Likewise, the number of downregulated genes in HEK-293T cells was also much lower (n = 5). Both types of T cells had nearly negligible responses to Lipofection (Figure 2E/F), with only three significantly upregulated genes in Jurkat (MT1E, MT1F, TMEM238) and a single upregulated gene in primary T cells (MT1H). No genes were significantly downregulated in either type of T cell following Lipofection.

### Validation of NGS with rt^2^PCR and ELISA

A select number of upregulated genes that were detected in PC-3 cells via mRNA-sequencing (Figure 2C) were also evaluated using rt^2^PCR and ELISA. Figure 3A shows that while there were some differences in the magnitude of upregulation observed with mRNA-sequencing (green bars) and rt^2^PCR (blue bars), 7 different genes (IFNB1, IFNL1/2/3, CXCL10/11, & CASP1) that were observed to be upregulated in mRNA-sequencing experiments were also found to be significantly upregulated in rt^2^PCR assays, thereby reinforcing the mRNA-sequencing results.

**Figure 3.**
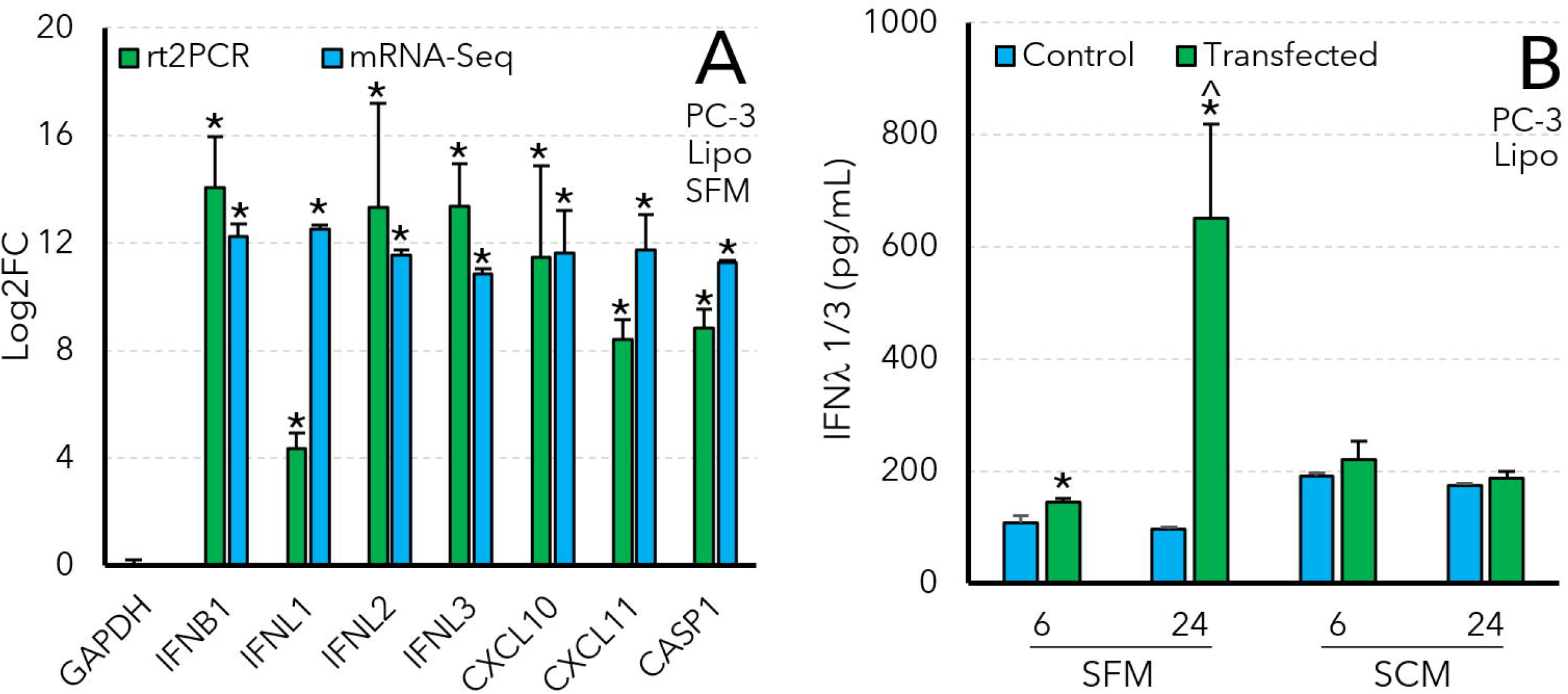
Validation of mRNA-sequencing results with rt^2^PCR (A) and ELISA (B). Asterisks (**\***) indicate significant increases in target levels following transfection relative to untransfected control cells, while carets (^) indicate significant increases between ELISA measurements at 6 and 24 hours after transfection.

Likewise, an increase in IFN*λ* 1/3 secretion was also verified with ELISA. It is worth noting that the antibody used in the ELISA assay binds to both IFN*λ* 1 and IFN*λ* 3, but significant increases in both IFN*λ*1 and IFN*λ*3 expression levels were detected in PC-3 cells at 6 and 24 hours after Lipofection in SFM (Figure 3B). IFN*λ* 1/3 levels were also significantly higher at 24 hours after Lipofection than 6 hours post-transfection. However, when PC-3 cells were Lipofected in serum-containing media (SCM) instead of SFM, there was no significant increase in IFN*λ* 1/3 expression levels between transfected and untransfected controls.

### Comparison of Transfection Methods in PC-3 Cells

The observation that IFN*λ* 1/3 levels did not increase following Lipofection of PC-3 cells in SCM (Figure 3) motivated us to further investigate the effects of serum on the transfection efficiency and transcriptome of PC-3 cells. In addition, we also investigated the effects of electroporation on the PC-3 transcriptome, since electroporation is a popular nonviral gene delivery method that can provide relatively high transfection efficiencies.

Indeed, Figure 4A shows that electroporation (in SCM) provided a much higher transfection efficiency (92 ± 2% GFP^+^ cells) in PC-3 cells at 24 hours post-electroporation. Likewise, the presence of serum during Lipofection also provided a slight, yet significant, increase in the transfection efficiency of the PC-3 cells (53 ± 4% GFP^+^ cells). In contrast, Lipofection of Jurkat and Primary T cells in SCM did not significantly increase transfection efficiency (data not shown).

**Figure 4:**
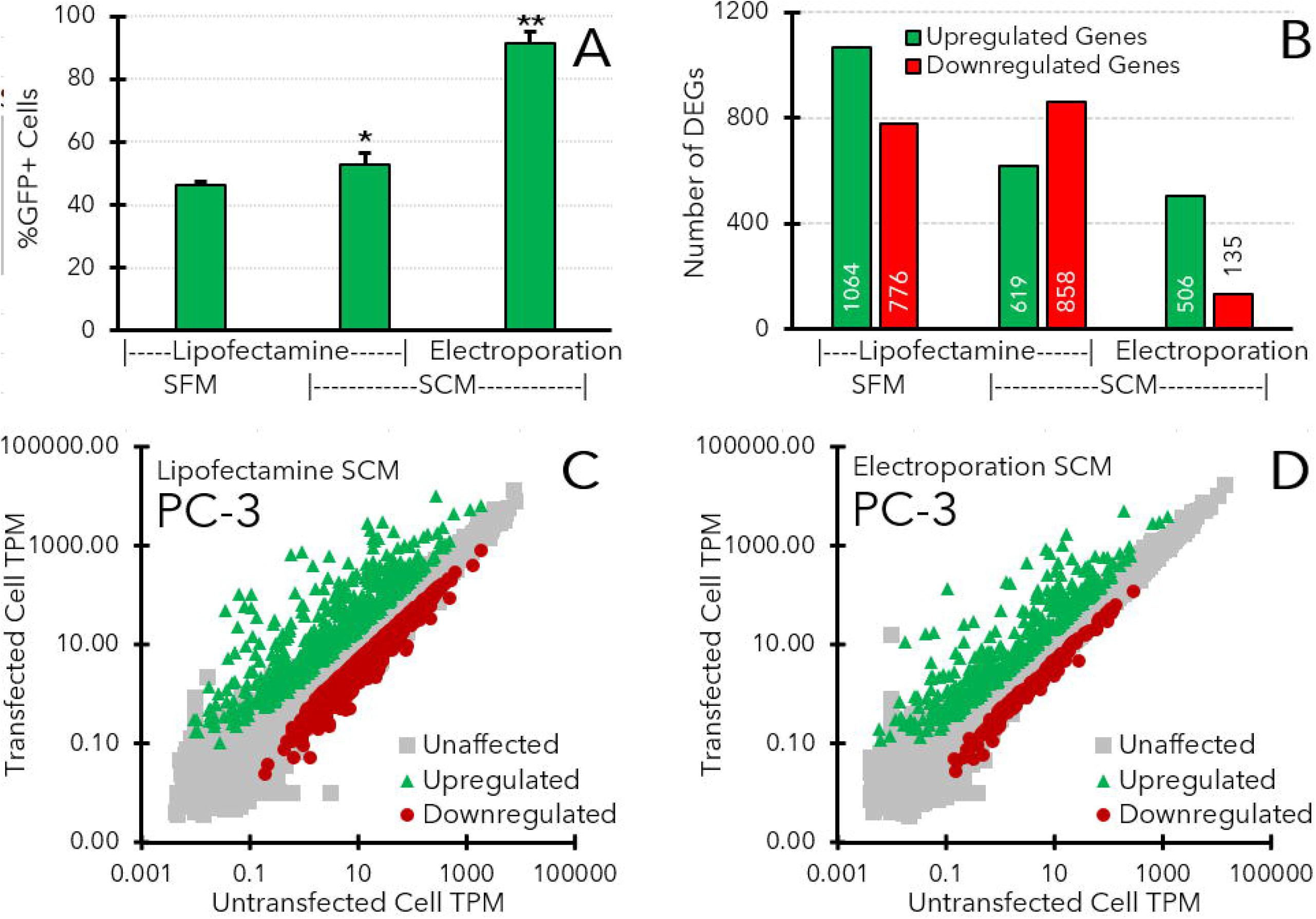
Effects of serum and electroporation on transfection efficiency and the PC-3 transcriptome. (A) Transfection efficiencies obtained with PC-3 cells using Lipofectamine in either SFM or SCM and Electroporation in SCM (blue bars, left axis). (B) A summary of the number of genes that were upregulated (green bars, right axis) or downregulated (red bars, right axis) following each type of transfection. Asterisks indicate significant differences in transfection efficiency (* = significantly higher than Lipofection in SFM, ** = significantly higher than Lipofection in SFM and SCM, p < 0.05 as determined by Student’s t-test). (B/C) Gene expression levels (TPM) for PC-3 cells that were Lipofected (B) or electroporated (C) in SCM. All TPM values are averaged from 3 separate transfections and corresponding mRNA-sequencing experiments. Green triangles indicate genes that were significantly upregulated in transfected cells, red circles indicate downregulated genes, and gray squares represent unaffected genes.

mRNA-sequencing revealed that the presence of serum during Lipofection of PC-3 cells decreased the number of genes that were significantly upregulated (n = 619, Figure 4B) compared to SFM (n = 1,057). Likewise, electroporation also resulted in a substantial decrease in the number of upregulated and downregulated genes (n = 533 and n = 138, Figure 4C) compared to Lipofection in both SFM and SCM. A complete list of the DEGs identified in these experiments is shown in a worksheet in the supplementary information.

Overall, the transfection efficiency of the PC-3 cells appeared to be inversely correlated with the number of DEGs. Electroporation provided the highest transfection efficiency and lowest number of DEGs, while Lipofection in SFM was associated with the highest number of DEGs and the lowest transfection efficiency.

### Overexpression of IFI16 in HEK-293T Cells

Analysis of the gene expression patterns in the different cell lines identified several genes (e.g., IFI16, IRF1, PSMB8, and PSMB9) with expression levels that appeared to be inversely correlated with the transfection efficiency of the host cell (Figure 5). For example, IFI16 was absent in HEK-293T cells, sharply upregulated in PC-3 cells, and constitutively expressed in both T cell lines. To test the hypothesis that genes like IFI16 might be interfering with transgene expression, IFI16 was transiently overexpressed from a plasmid in HEK-293T cells and then subsequently transfected with pEF-GFP 24 hours later to determine if IFI16 might inhibit GFP expression in HEK-293T cells. Indeed, Figure 6 shows that the HEK-293T cells that were only transfected with pEF-GFP achieved a high transfection efficiency (80.3%), while cells that were co-transfected with pIFI16 and pEF-GFP obtained at a significantly lower transfection efficiency (33.2%). To ensure that this lower transfection efficiency was not solely due to the dual transfections, additional samples were co-transfected with the luciferase expression plasmid pGL4.50 and pEF-GFP using the same protocol. Cells in this category did exhibit a significantly lower transfection efficiency (65.5%) than the cells transfected with only pEF-GFP, but a Wilcoxon Rank Sum test showed a significant decrease in transfection efficiency between the pGL4.50 group and the IFI16 group (p = 2 ⨯ 10^−5^).

**Figure 5.**
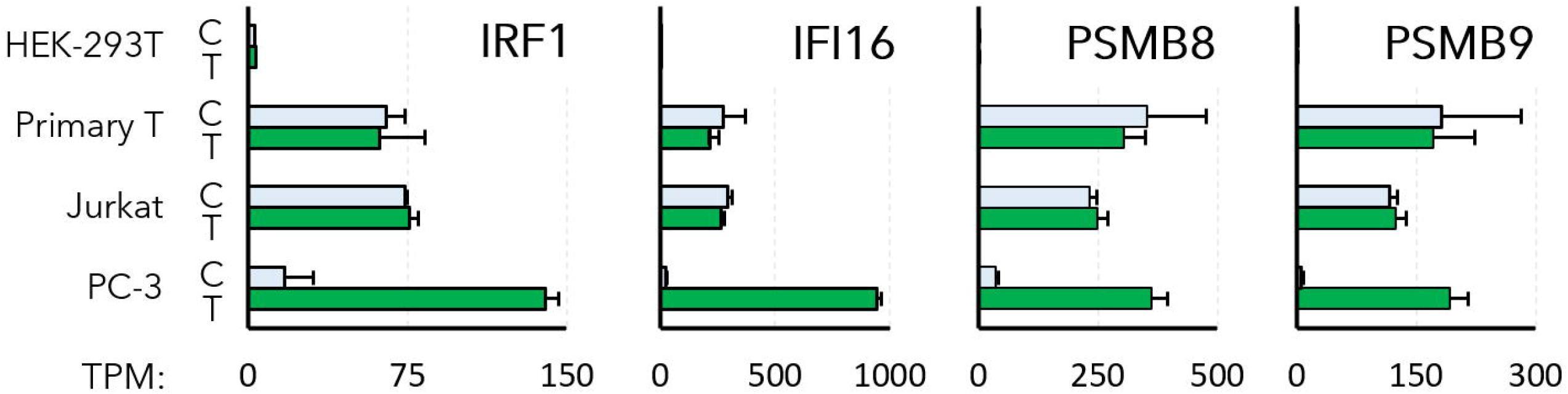
Expression levels (TPM) of five representative cytokine-stimulated genes (CSGs) that were observed to be absent or expressed at low levels in HEK-293T cells, upregulated during Lipofection in PC-3 cells, and constitutively expressed at relatively high levels in Jurkat and Primary T cell lines.

**Figure 6.**
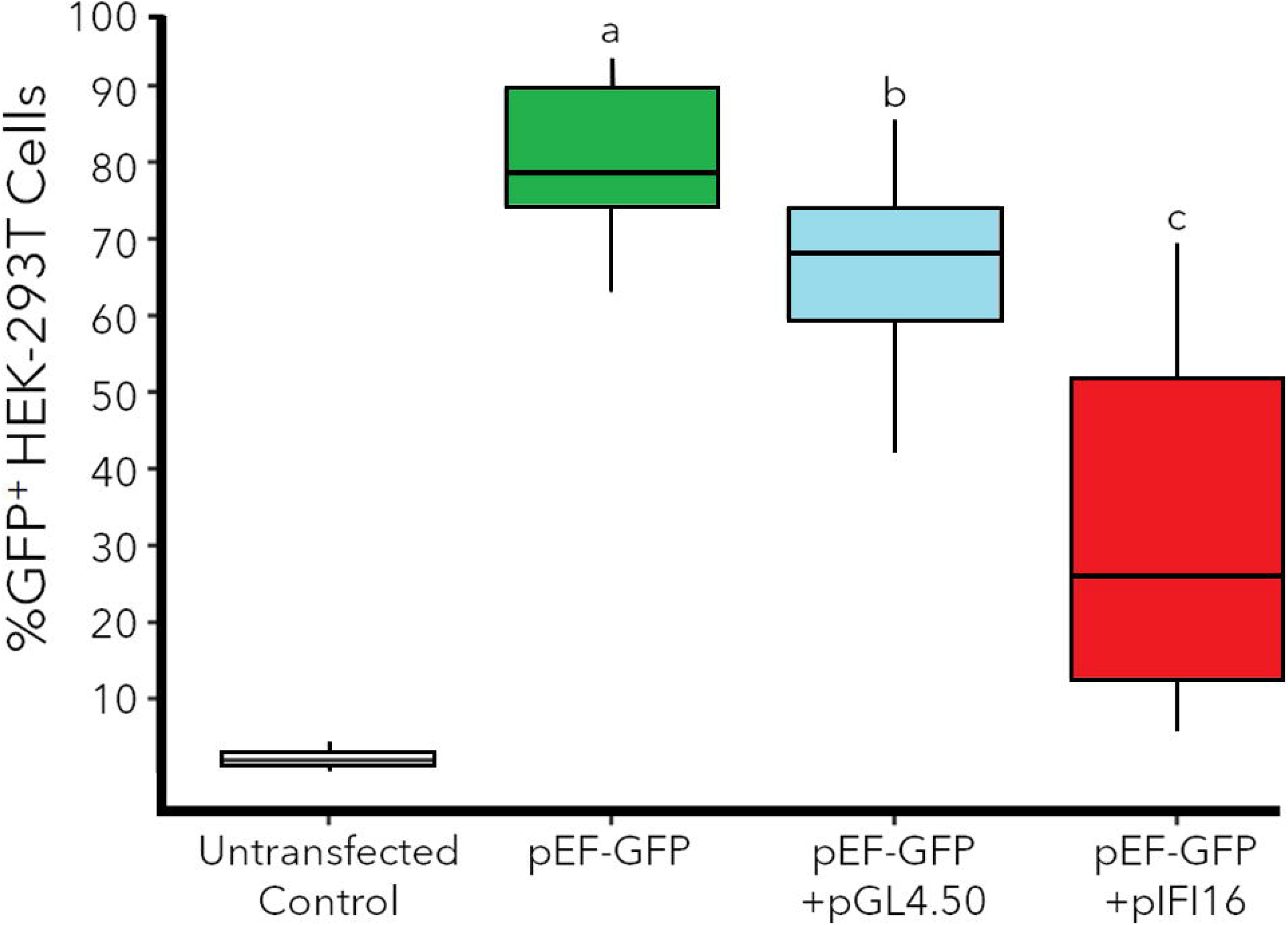
Effects of IFI16 overexpression on pEF-GFP transfection efficiency (%GFP^+^ cells) in HEK-293T cells. Untransfected negative control cells were not transfected, while positive control cells were only transfected with pEF-GFP (green box). The blue box represents cells that were transfected with the luciferase expression plasmid pGL4.50 one day prior to transfection with pEF-GFP, while the red box represents cells that were transfected with the IFI16 expression plasmid pIFI16-FL one day before transfection. Horizontal lines within boxes represent the means for each group, while boxes show the interquartile range, and the entire dataset is contained within the whiskers. Letters (a, b, c) indicate samples with significantly different transfection efficiencies (n = 18 for each sample, p < 0.05 using a Wilcoxon Rank Sum test).

## DISCUSSION

### Differences in DNA Sensing Pathways

One of the most drastic differences between the transcriptomic profiles shown in Figure 2 was a decrease in the number of upregulated genes between PC-3 cells (n = 1,057) and the HEK-293T (n = 142) cells. Furthermore, it is important to note that while several inflammatory cytokines (e.g., IFNβ, IFN*λ*, and IL-6) and CSGs were upregulated in PC-3 cells, no cytokines and CSGs were detected in HEK-293T cells (Table S2). This dampened innate immune response suggests that HEK-293T cells may be missing one or more components of the DNA sensing pathways that drive the potent innate immune response in PC-3 cells.

Indeed, multiple components of the CpG-DNA sensing pathways were not detected in HEK-293T cells (Table 1), including the endosomal CpG-DNA sensor TLR9. TLR9 was also absent in PC-3, Jurkat, and primary T cells, but the cytosolic CpG-DNA sensors DHX9 and DHX36 were detected in all the cell lines.^30^ However, the downstream adaptor protein MyD88 that is required for signaling by DHX9 and DHX36 (Figure 1) was also expressed at a relatively low level in HEK-293T cells (TPM = 2.0 vs. TPM = 298.4 in PC-3 cells). Therefore, while CpG-DNA sensing by DHX9 and DHX36 might contribute to the innate immune response to pDNA in PC-3 cells, the lack of MyD88 in HEK-293T cells may prevent them from sensing foreign CpG DNA.

**Table 1.**
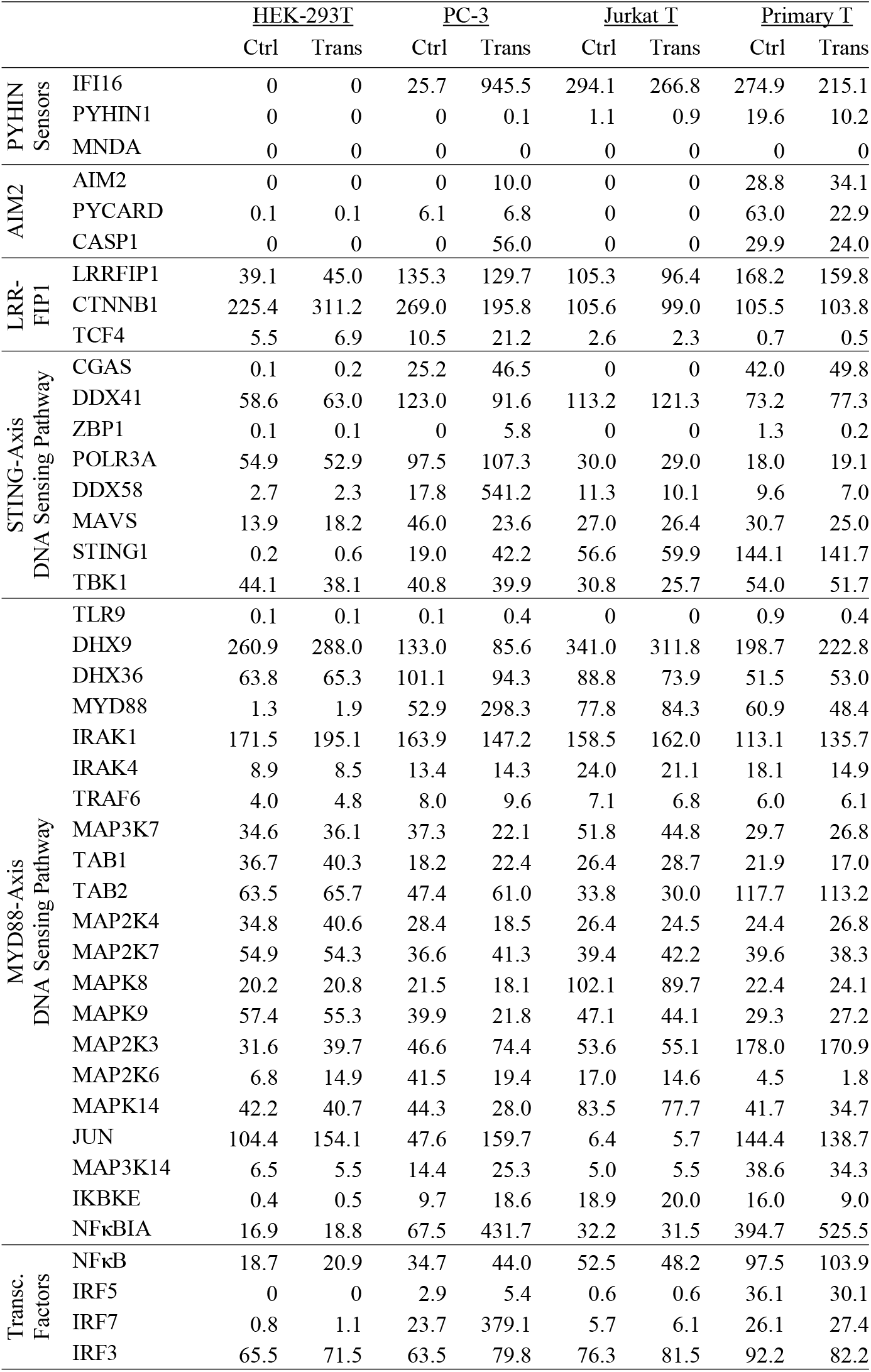
Expression levels (TPM) of genes involved in DNA sensing pathways in each cell line.

The STING axis is an alternative cytosolic DNA sensing pathway that may also drive the innate immune response to pDNA in PC-3 cells. PC-3 cells express each component of the STING-mediated DNA sensing pathway (Table 1), which is induced when IFI16 or cGAS bind to cytosolic dsDNA and then signal through STING to activate TBK-1, IRF3, and NF-κB (Figure 1). However, several important components of this pathway were either absent or expressed at relatively low levels in HEK-293T cells. For example, IFI16 was detected in PC-3 cells and upregulated 37-fold following Lipofection, but it was absent in HEK-293T cells. Three more components of the STING pathway (cGAS, STING, and IKBKE) were also upregulated in the transfected PC-3 cells but expressed at nearly negligible levels (TPM = 0.1-0.6) in HEK-293T cells. Altogether, these data show that while PC-3 cells can sense pDNA with multiple redundant pathways and mount a potent innate immune response that induces hundreds of cytokines and CSGs, HEK-293T cells lack essential proteins at bottlenecks in DNA sensing pathways (e.g., MyD88 and STING), which may explain the lack of cytokine and CSG expression shown in Tables S2 and S3, respectively.

It is worth noting that 142 genes were upregulated in the transfected HEK-293T cells, but those genes do not have any known functions that could potentially inhibit transgene expression. Instead, it is possible that those genes may have been upregulated in response to the stress or toxicity caused by the presence of Lipofectamine.

Many other groups have also observed the high transfection efficiency exhibited by HEK-293T cells, which has led to their widespread adoption in industry and academia.^31–33^ The high transfection efficiency and lack of CSG expression in HEK-293T cells may be due to the SV40 large T antigen, which was integrated into the parental HEK-293 cell genome to create the HEK-293T cell line.^34^ The SV40 large T antigen has previously been shown to improve the replication efficiency of DNA viruses^35^ and inhibit the induction of interferon expression by the cGAS-STING signaling pathway in other cell lines.^33^ Likewise, proteins expressed by the human papilloma virus (HPV) have also been shown to inhibit STING,^36^ while the absence of STING in hepatocytes has been correlated with a lack of cytokine expression during hepatitis B virus (HBV) infection.^37^ In regards to nonviral gene delivery, the inhibition of STING with small molecule inhibitors (C176, C178) has been shown to enhance transgene expression by up to 3-fold in a variety of cell lines, including primary T cells.^38^ These previous studies and our observations collectively emphasize the importance of the STING DNA sensing pathway and suggest that targeting STING for inhibition may be an effective strategy to improve the potency of gene therapy treatments.

### Constitutive Expression of CSGs in T cells

Another interesting observation is that the Jurkat and primary CD3^+^ T cells showed almost no response to Lipofection, except for the upregulation of a few metallothioneins (MT1H, MT1E, and MT1F). Metallothioneins have previously been shown to restrict bacterial and viral replication, but it is unclear how they may interfere with nonviral gene delivery or expression^39,40^.

In contrast to the PC-3 cells, no interferons or CSGs were upregulated following Lipofection in either T cell line (Table S2). It is worth noting that some chemokines (CXCL8/10/13) and TNFα were detected in the primary T cells, but each of these targets was detected in both the transfected and untransfected cells. Therefore, it is more likely that these cytokines were induced by the activation of the primary T cells with IL-2 rather than the transfection of plasmid DNA.

The lack of interferon and CSG upregulation following Lipofection of the T cells is somewhat surprising, since transcripts for all the requisite components of multiple DNA sensing pathways were detected in the T cells (e.g., IFI16, STING, TBK-1, and IRF3). Therefore, the T cells should be able to induce expression of cytokines, but interleukins and interferons like IFN*λ*1 were not expressed in the T cells following transfection (Table S2). Similar observations were made in a previous study that also detected IFI16 and TBK-1 expression in T cells, but no interferon expression.^41^ Therefore, it appears that T cells may either lack an unknown component that is required for cytokine induction or the interferon genes may be epigenetically silenced.^42^

Although interferons and CSGs were not upregulated in T cells after Lipofection, several CSGs with established anti-viral activities were constitutively expressed in both transfected and untransfected T cells. Furthermore, many of the CSGs that were expressed in the T cells were completely absent in the easily transfected HEK-293T cells and significantly upregulated in the PC-3 cells that had a moderate transfection efficiency. This inverse correlation between the expression levels of these CSGs and the transfection efficiency of each host cell line suggests that these CSGs may inhibit transgene uptake or expression. A complete list of the CSGs which follow this trend (absent in HEK-293T, upregulated in PC-3, and constitutively expressed in T cells) is shown in Table S3, while the TPMs of some representative CSGs in each cell line are shown in Figure 5.

While Table S3 consists of a wide variety of 54 different CSGs, the four genes highlighted in Figure 5 are the most likely to inhibit transgene expression. For example, the constitutive expression of CSGs in the absence of interferons may be driven by the transcription factor IRF1, which was upregulated 8-fold in PC-3 cells after Lipofection and detected in all T cell samples (TPM = 62-76). Unlike other interferon regulatory factors (IRFs), IRF1 does not require phosphorylation to activate its target genes. Therefore, IRF1 may continuously drive high level expression of CSGs in T cells.^43^

IFI16 is another important anti-viral restriction factor. As previously mentioned, IFI16 is a cytosolic DNA sensor, but recent studies have revealed additional anti-viral functions for IFI16.^44^ For example, IFI16 can repress viral genes by directly binding viral promoters to inhibit transcription.^27,45,46^ Some viruses have adapted to this problem by expressing inhibitory proteins (e.g., ICP0 from HSV-1) that induce the degradation of IFI16 and prevent its transcriptional repression.^25^ Therefore, while our results show that IFI16 is unable to trigger the expression of interferons and other cytokines in T cells, IFI16 may instead repress the transcription of both viral and nonviral transgenes by binding to their upstream promoters. Additionally, IFI16 is known to form an inflammasome complex with PYCARD, Caspase 1/8, and Gasdermin D upon binding to foreign DNA which can lead to inflammation and cell death. Indeed, pyroptosis has been observed during abortive HIV infection of T cells.^47,48^

Several of the CSGs listed in Table S3 have very well-established roles in the adaptive and innate immune responses to viral infection, but it is unclear how they might inhibit the expression of nonviral transgenes. For example, PSMB8 and PSMB9 are two closely related CSGs that were upregulated in PC-3 cells and expressed at similarly high levels in both T cell lines. PSMB8 and PSMB9 have been shown to regulate transcription changes during the innate immune response, but they are more widely known for their role in the assembly and function of the immunoproteasome, which selectively degrades foreign proteins into peptide antigens that are then displayed on MHC-1.^49–51^ This process is essential to the recruitment of cytotoxic CD8^+^ T cells to virus-infected cells *in vivo*, but it is not yet known if the immunoproteasome may specifically interfere with transgene expression.

### Effects of Serum and Electroporation on the PC-3 Transcriptome

Our initial transfections were conducted in serum-free media (SFM) to avoid any potentially confounding effects on the transcriptome from components in the serum.^52–54^ However, serum components like albumin have also been reported to enhance Lipofection efficiency.^55,56^ Likewise, several studies have shown that electroporation can provide relatively high transfection efficiencies, even in cell lines that are relatively difficult to transfect.^21,22^ As shown in Figure 4A, electroporation in SCM also yielded significantly higher transfection efficiencies in PC-3 cells than Lipofection in SFM.

While it is possible that electroporation achieves a higher transfection efficiency by simply delivering more of the transgene to the cell, there are some intriguing differences in gene expression that were observed between the Lipofected and electroporated cells. For example, the TPMs of several DNA sensors, cytokines, and CSGs seemed to be inversely correlated with the transfection efficiencies of the different types of transfected PC-3 cells. Table 2 shows some of the more noteworthy genes that were highly upregulated after Lipofection with SFM but expressed at significantly lower levels in electroporated PC-3 cells.

**Table 2.**
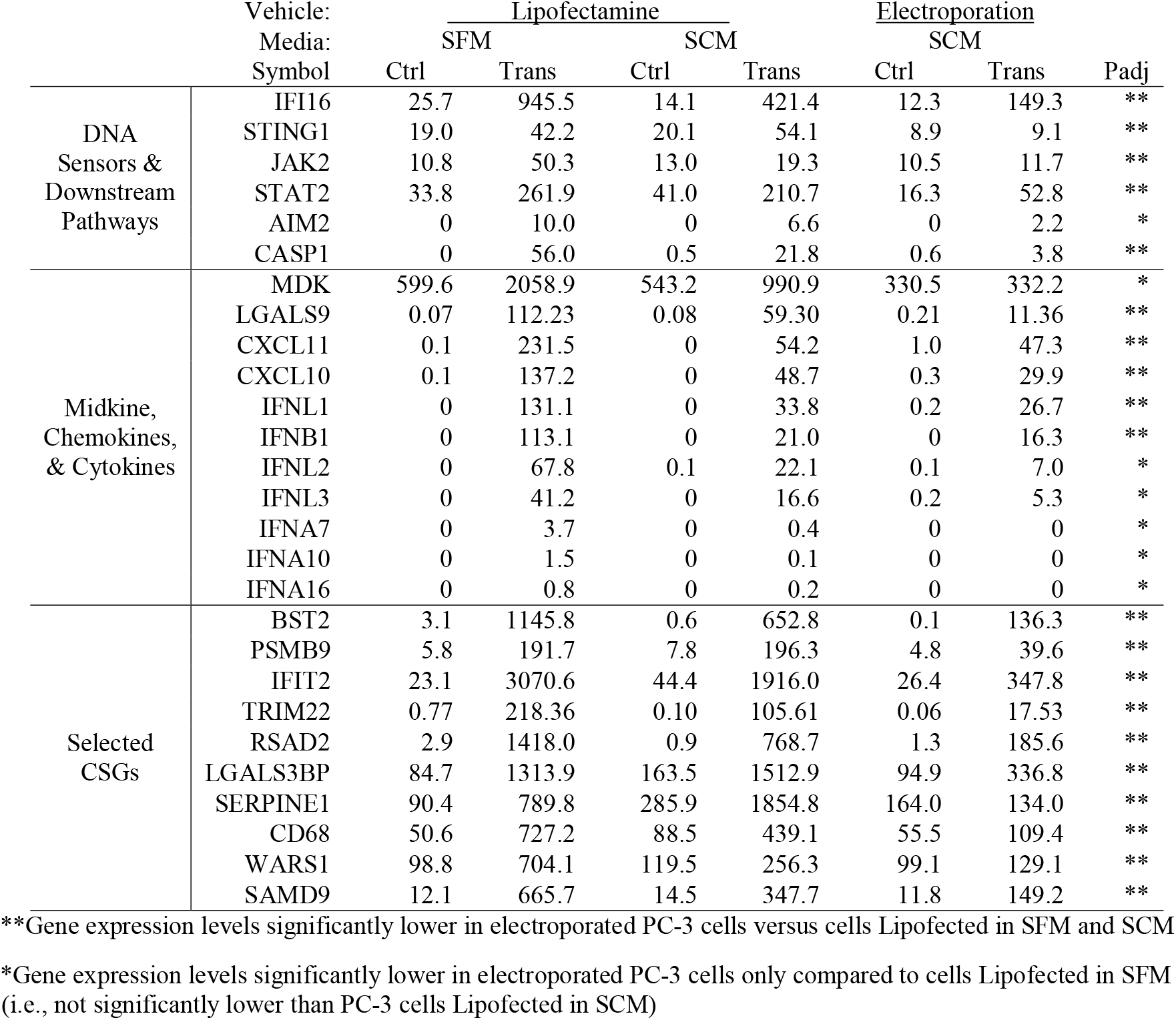
Genes expressed at significantly lower levels in electroporated PC-3 cells vs. Lipofected PC-3 cells

Several components of the STING axis of DNA sensing were expressed at significantly lower levels in the electroporated PC-3 cells. IFI16 emerged again as a notable example of this trend since it was highly upregulated (TPM = 945.5) following Lipofection in SFM but expressed at significantly lower levels during Lipofection in SCM (TPM = 421.4) and electroporation (TPM = 149.3). Other important genes that were expressed at lower levels in electroporated PC-3 cells include STING, mediators of cytokine signaling (JAK2 and STAT2), and components of the inflammasome pathway (AIM2 and Caspase I) that induce inflammation or apoptosis in response to cytosolic DNA.^22^ Altogether, these observations show that DNA sensing pathways may be less active in electroporated PC-3 cells, which could explain the lower number of cytokines and CSGs observed in those samples.

Indeed, multiple chemokines and interferons that were highly expressed in PC-3 cells following Lipofection in SFM were expressed at much lower levels in the electroporated cells. Table 2 shows that all the interferons (IFNα, IFNβ, and IFN*λ*s) were expressed at lower levels following electroporation, along with the chemokines CXCL 10 and 11. Another intriguing observation is that the expression of two other secreted proteins - midkine (MDK) and LGALS9 – decreased slightly in the presence of serum and to much lower levels (6 to 10-fold lower) during electroporation. LGALS9 and MDK have previously been shown to inhibit the initial binding of viral capsids and cationic lipoplexes to cell membranes by blocking heparan sulfate proteoglycans (HSPGs), syndecans (SDCs), and other cell surface receptors that are crucial for gene delivery.^57–61^ Therefore, it is possible that the lower transfection efficiencies observed in PC-3 cells cultured in SFM may be due to inhibition of lipoplex endocytosis by MDK or LGALS9.

The importance of HSPGs and SDCs in gene delivery is also highlighted by the observation that these genes were expressed at much lower levels in T cells than PC-3 and HEK-293T and PC-3 cells (see Table S4). Previous studies have also shown this lack of HSPG expression in T cells,^77^ which provides another explanation for their relatively low transfection efficiencies.

Finally, many other CSGs that were expressed at lower levels after electroporation than Lipofection in SFM are listed in Table 2 and the worksheet in the supplementary information. The functions of many of these genes have not yet been revealed, but there are some CSGs with known functions that might inhibit transgene expression. For example, much like IFI16, TRIM22 can specifically bind transgene promoters to exclude other activating transcription factors (e.g., SP-1).^46,62^ Alternatively, TRIM22 has also been shown to ubiquitinate viral proteins to target them for degradation.^63^

### Inhibition of Transgene Expression by IFI16

While there are many CSGs that could potentially inhibit transgene expression, the strongest inverse correlation between CSG expression levels and host cell transfection efficiencies was observed for IFI16. Indeed, the lack of IFI16 in HEK-293T cells may explain their lack of cytokine expression and high transgene expression levels, while the high levels of IFI16 expression in T cells and PC-3 cells may decrease transgene expression. However, it is important to note that IFI16 expression levels were much lower in PC-3 cells following electroporation, which may explain the much higher electroporation efficiencies observed in PC-3 cells.

Figure 6 also shows that overexpression of IFI16 in HEK-239T cells subsequently decreases GFP expression. This decrease in transfection efficiency is probably not due to the DNA sensing activity of IFI16, since HEK-293T cells lack STING and TBK-1. Instead, it is more likely that IFI16 directly binds to the transgene to repress its transcription, as previously described for other viral transgenes.^27,45,46^

Overall, our analysis of the transcriptomes of several types of transfected cells emphasizes the importance of DNA sensing pathways and specific CSGs (e.g., IFI16, IRF1, et al.) in the innate immune response to transgene delivery. Our findings contribute to a growing body of literature that indicate the STING axis is particularly important for DNA sensing and subsequent cytokine expression, since the expression levels of IFI16 and STING are inversely correlated with transfection efficiencies in multiple cell lines and transfection methods. Our results also show that while PC-3 cells exhibit a potent innate immune response that limit their Lipofection efficiency in serum-free media, it is possible to dampen the innate immune response and increase transfection efficiency by adding serum or using electroporation to deliver pDNA.

Finally, our mRNA-sequencing experiments have identified multiple possible reasons for the notoriously low Lipofection efficiencies observed for T cells in this study and previous studies, including a lack of HSPGs and the constitutive expression of repressive CSGs like IFI16 by IRF1. Several other studies have demonstrated that IFI16 can inhibit transgene delivery and expression, but additional knockout studies will be necessary to confirm the potential roles of the other CSGs in transgene expression. However, after transgene repressors have been identified, short interfering RNAs (siRNAs) or small molecule inhibitors could be developed to inhibit these targets and potentially improve the potency of future viral and future nonviral gene therapies.

## MATERIALS AND METHODS

### Reagents

Lipofectamine LTX was purchased from ThermoFisher Scientific (#15338100). The GFP and IFI16 expression plasmids were purchased from Addgene (Plasmids 11154 and 35051, respectively. Watertown, MA), while the luciferase expression plasmid pGL4.50 was purchased from Promega (cat# E1310, Madison, WI).

### Cell Lines

Human PC-3 prostate cancer (Cat# CRL-1573), Jurkat T lymphoma (TIB-152), and HEK-293T embryonic kidney (CRL-11268) cells were purchased from ATCC, while donated primary CD3^+^ samples of T cells from 3 different donors were purchased from Cellero (formerly known as Astarte Bio). PC-3, Jurkat, and HEK-293T cells were cultured in serum-containing RPMI-1640 media that was supplemented with 10% fetal bovine serum, except during the 24-hour period following a transfection (unless otherwise noted). In contrast, Primary T cells were cultured in serum-free X-VIVO15 media that was supplemented with 0.5 ng/mL IL-2 and anti-CD3/28 Dynabeads in a 1:1 cell:bead ratio. Dynabeads were replaced weekly, while IL-2 was added to the media every 2-3 days.

### Transfections

In preparation for mRNA sequencing, each cell line was transfected using specific conditions that maximized transfection efficiency (data not shown). Adherent cell lines (PC-3 and HEK-293T) were grown to 50-70% confluency in T-75 flasks (∼5M cells/flask) and then transfected with lipoplexes that were prepared by mixing pEF-GFP (4.5 ug) with Lipofectamine LTX (9 uL) and PLUS reagent (4.5 uL) in ∼200 uL of OptiMEM media.

Jurkat T cells were seeded into T-25 flasks (2.5⨯10^6^ cells) and then transfected with lipoplexes that were prepared by mixing pEF-GFP (13.2 ug) with Lipofectamine LTX (36.2 uL) and PLUS reagent (13.2 uL) in ∼200 uL of OptiMEM media. Smaller cultures of primary CD3^+^ T cells were seeded at a density of 200,000 cells/well and then transfected with lipoplexes that were prepared by mixing 1 ug pEF-GFP/well, 2.75 uL Lipofectamine LTX/well, and 1 uL PLUS reagent/well.

With the exception of the Lipofection and electroporation experiments conducted with serum-containing media (SCM) in PC-3 cells, the SCM was removed from the cells in all other experiments and replaced with serum-free media (SFM) before the lipoplexes were added to each of the cell lines. Cells were then subsequently incubated for 24 hours at 37°C in SFM. Finally, a Millipore Guava flow cytometer was used to quantify transfection efficiency (%GFP cells) and transgene expression (mean GFP) prior to RNA isolation.

### IFI16 Overexpression in HEK-293T Cells

HEK-293T cells were seeded into 24-well plates at 25,000 cells/well in SCM and incubated at 37°C on Day 1. The wells on each plate were then divided into 4 groups – a negative control group that was never transfected, a positive control group that was transfected once with pEF-GFP on Day 3, and two other groups that were transfected with either pGL4.50 (a luciferase expression plasmid) or pIFI16-IL (IFI16 expression plasmid) on Day 2 and then transfected again with pEF-GFP on Day 3. In each transfection, plasmids were administered at 1µg DNA/well along with 1µL/well Lipofectamine LTX and 0.5µL/well PLUS reagent. Plates were incubated for an additional 48 hours at 37°C and transfection efficiency (%GFP cells) was measured using a Millipore Guava flow cytometer on Day 5.

### Electroporation

PC-3 cells were trypsinized, centrifuged at 1,000 RPM for 4.5 minutes, and then washed once with PBS before centrifuging the cells again and resuspending them in PBS again. Plasmid DNA (7.5 μg pEF-GFP) and 2.5 ⨯ 10^6^ PC-3 cells were then mixed in a total volume of 100 μL in sterile 2 mm-gap electroporation cuvettes (in that order). A Bio-Rad Gene Pulser Xcell electroporation system with the CE module was then used to briefly electroporate the cells (110V, 25 ms, single square wave pulse). The electroporated cells were then immediately plated in preheated RPMI media with serum and incubated at 37°C until needed.

### mRNA Sequencing

Total RNA samples were extracted from cell samples with a Qiagen RNEasy kit. The RNA samples (2 μg total RNA) were then submitted to either Genewiz (Jurkat samples) or the Beijing Genomics Institute (PC-3, HEK-293T, and Primary T cell samples) for library preparation and mRNA-sequencing. Specifically, mRNA was isolated from high quality total RNA samples with RIN > 9 and 28S:18S > 1 (measured with an Agilent 2100 Bioanalyzer) using poly-T oligonucleotide beads. The mRNAs were then cleaved into smaller fragments that were subsequently reverse transcribed by DNA Polymerase I into cDNA using random N6 primers. Adapters were then ligated onto the cDNAs and the resulting libraries were sequenced using either a BGI-500 sequencer (BGI) or an Illumina HiSeq (Genewiz). Low quality reads were then filtered to produce clean reads that were mapped to the human genome/transcriptome using HISAT, Bowtie2, and RSEM to calculate gene expression values/counts. Counts were then analyzed using DESeq2 in RStudio with independent filtering turned off (see Figure S2) to identify differentially expressed genes (DEGs), which were defined as genes with adjusted p-values (p_adj_) less than 0.05 and at least a 2-fold change in TPM following transfection.

Transcripts per million (TPM) values were calculated by dividing the number of counts for each gene by each gene’s length to obtain reads per kilobase (RPK) values. The RPK for each gene was then divided by the sum of the RPK values measured for all genes (and multiplied by one million) to obtain TPM values. The complete mRNA-seq data (fastq files and a spreadsheet of TPM values) from these experiments are available at the NCBI GEO repository (GEO Accession #GSE166630).

### NGS Validation with rt^2^PCR

The same RNA samples that were used for mRNA-sequencing were also analyzed with rt^2^PCR. First, mRNA was reverse transcribed using the SuperScript IV VILO™ rt^2^PCR master mix with ezDNase™ (ThermoFisher #11766050) according to the manufacturer’s protocol. The resulting cDNA was then quantified using a SYBR green qPCR master mix (ThermoFisher # 4472942) with a QuantStudio 3 qPCR instrument (ThermoFisher). All measurements were repeated in triplicate and the housekeeping gene glyceraldehyde-3-phosphate dehydrogenase (GAPDH) was used as a reference gene/control. The primer sequences that were used are shown in Table S1.

### Cytokine Quantification with ELISA

Samples of the supernatant media were taken from PC-3 cultures at 6 and 24 hours after transfection with Lipofectamine and stored at -72^**o**^C until needed. DuoSet ELISA kits were then used to quantify IFNL1/3 (Biotechne #DY1598B-05) levels according to the manufacturer’s protocol.

## ACKNOWLEDGEMENTS

This work was supported by the U.S. National Science Foundation (Grants #1645225 and #1651837). The pEF-GFP plasmid used for transfections was a gift from Connie Cepko (Addgene plasmid # 11154) while the pIFI16-FL plasmid was a gift from Cheryl Arrowsmith (Addgene plasmid # 35051).

## AUTHOR CONTRIBUTIONS

Jacob Elmer: Conceptualization, Methodology, Supervision, Validation, Data Curation, Writing, Project Administration, & Funding Acquisition. Eric Warga: Investigation, Formal Analysis, Validation, & Writing. Matthew Tucker: Investigation. Emily Harris: Investigation.

