## Supplementary material for "Transcriptomic Analysis of the Innate Immune Response to *in vitro* Transfection of Plasmid DNA": Tables

**Table 1.** Expression levels (TPM) of genes involved in DNA sensing pathways in each cell line.

|  |  | HEK-293T | | PC-3 | | Jurkat T | | Primary T | |
| --- | --- | --- | --- | --- | --- | --- | --- | --- | --- |
|  |  | Ctrl | Trans | Ctrl | Trans | Ctrl | Trans | Ctrl | Trans |
| PYHIN Sensors | IFI16 | 0 | 0 | 25.7 | 945.5 | 294.1 | 266.8 | 274.9 | 215.1 |
|  | PYHIN1 | 0 | 0 | 0 | 0.1 | 1.1 | 0.9 | 19.6 | 10.2 |
|  | MNDA | 0 | 0 | 0 | 0 | 0 | 0 | 0 | 0 |
| AIM2 | AIM2 | 0 | 0 | 0 | 10.0 | 0 | 0 | 28.8 | 34.1 |
|  | PYCARD | 0.1 | 0.1 | 6.1 | 6.8 | 0 | 0 | 63.0 | 22.9 |
|  | CASP1 | 0 | 0 | 0 | 56.0 | 0 | 0 | 29.9 | 24.0 |
| LRR-FIP1 | LRRFIP1 | 39.1 | 45.0 | 135.3 | 129.7 | 105.3 | 96.4 | 168.2 | 159.8 |
|  | CTNNB1 | 225.4 | 311.2 | 269.0 | 195.8 | 105.6 | 99.0 | 105.5 | 103.8 |
|  | TCF4 | 5.5 | 6.9 | 10.5 | 21.2 | 2.6 | 2.3 | 0.7 | 0.5 |
| STING-Axis  DNA Sensing Pathway | CGAS | 0.1 | 0.2 | 25.2 | 46.5 | 0 | 0 | 42.0 | 49.8 |
|  | DDX41 | 58.6 | 63.0 | 123.0 | 91.6 | 113.2 | 121.3 | 73.2 | 77.3 |
|  | ZBP1 | 0.1 | 0.1 | 0 | 5.8 | 0 | 0 | 1.3 | 0.2 |
|  | POLR3A | 54.9 | 52.9 | 97.5 | 107.3 | 30.0 | 29.0 | 18.0 | 19.1 |
|  | DDX58 | 2.7 | 2.3 | 17.8 | 541.2 | 11.3 | 10.1 | 9.6 | 7.0 |
|  | MAVS | 13.9 | 18.2 | 46.0 | 23.6 | 27.0 | 26.4 | 30.7 | 25.0 |
|  | STING1 | 0.2 | 0.6 | 19.0 | 42.2 | 56.6 | 59.9 | 144.1 | 141.7 |
|  | TBK1 | 44.1 | 38.1 | 40.8 | 39.9 | 30.8 | 25.7 | 54.0 | 51.7 |
| MYD88-Axis  DNA Sensing Pathway | TLR9 | 0.1 | 0.1 | 0.1 | 0.4 | 0 | 0 | 0.9 | 0.4 |
|  | DHX9 | 260.9 | 288.0 | 133.0 | 85.6 | 341.0 | 311.8 | 198.7 | 222.8 |
|  | DHX36 | 63.8 | 65.3 | 101.1 | 94.3 | 88.8 | 73.9 | 51.5 | 53.0 |
|  | MYD88 | 1.3 | 1.9 | 52.9 | 298.3 | 77.8 | 84.3 | 60.9 | 48.4 |
|  | IRAK1 | 171.5 | 195.1 | 163.9 | 147.2 | 158.5 | 162.0 | 113.1 | 135.7 |
|  | IRAK4 | 8.9 | 8.5 | 13.4 | 14.3 | 24.0 | 21.1 | 18.1 | 14.9 |
|  | TRAF6 | 4.0 | 4.8 | 8.0 | 9.6 | 7.1 | 6.8 | 6.0 | 6.1 |
|  | MAP3K7 | 34.6 | 36.1 | 37.3 | 22.1 | 51.8 | 44.8 | 29.7 | 26.8 |
|  | TAB1 | 36.7 | 40.3 | 18.2 | 22.4 | 26.4 | 28.7 | 21.9 | 17.0 |
|  | TAB2 | 63.5 | 65.7 | 47.4 | 61.0 | 33.8 | 30.0 | 117.7 | 113.2 |
|  | MAP2K4 | 34.8 | 40.6 | 28.4 | 18.5 | 26.4 | 24.5 | 24.4 | 26.8 |
|  | MAP2K7 | 54.9 | 54.3 | 36.6 | 41.3 | 39.4 | 42.2 | 39.6 | 38.3 |
|  | MAPK8 | 20.2 | 20.8 | 21.5 | 18.1 | 102.1 | 89.7 | 22.4 | 24.1 |
|  | MAPK9 | 57.4 | 55.3 | 39.9 | 21.8 | 47.1 | 44.1 | 29.3 | 27.2 |
|  | MAP2K3 | 31.6 | 39.7 | 46.6 | 74.4 | 53.6 | 55.1 | 178.0 | 170.9 |
|  | MAP2K6 | 6.8 | 14.9 | 41.5 | 19.4 | 17.0 | 14.6 | 4.5 | 1.8 |
|  | MAPK14 | 42.2 | 40.7 | 44.3 | 28.0 | 83.5 | 77.7 | 41.7 | 34.7 |
|  | JUN | 104.4 | 154.1 | 47.6 | 159.7 | 6.4 | 5.7 | 144.4 | 138.7 |
|  | MAP3K14 | 6.5 | 5.5 | 14.4 | 25.3 | 5.0 | 5.5 | 38.6 | 34.3 |
|  | IKBKE | 0.4 | 0.5 | 9.7 | 18.6 | 18.9 | 20.0 | 16.0 | 9.0 |
|  | NFκBIA | 16.9 | 18.8 | 67.5 | 431.7 | 32.2 | 31.5 | 394.7 | 525.5 |
| Transc. Factors | NFκB | 18.7 | 20.9 | 34.7 | 44.0 | 52.5 | 48.2 | 97.5 | 103.9 |
|  | IRF5 | 0 | 0 | 2.9 | 5.4 | 0.6 | 0.6 | 36.1 | 30.1 |
|  | IRF7 | 0.8 | 1.1 | 23.7 | 379.1 | 5.7 | 6.1 | 26.1 | 27.4 |
|  | IRF3 | 65.5 | 71.5 | 63.5 | 79.8 | 76.3 | 81.5 | 92.2 | 82.2 |

**Table 2** – Genes expressed at significantly lower levels in electroporated PC-3 cells vs. Lipofected PC-3 cells

|  | Vehicle: | Lipofectamine _ | | | | Electroporation | |  |
| --- | --- | --- | --- | --- | --- | --- | --- | --- |
|  | Media:  Symbol | SFM | | SCM | | SCM | |  |
|  |  | Ctrl | Trans | Ctrl | Trans | Ctrl | Trans | Padj |
| DNA Sensors & Downstream Pathways | IFI16 | 25.7 | 945.5 | 14.1 | 421.4 | 12.3 | 149.3 | ** |
|  | STING1 | 19.0 | 42.2 | 20.1 | 54.1 | 8.9 | 9.1 | ** |
|  | JAK2 | 10.8 | 50.3 | 13.0 | 19.3 | 10.5 | 11.7 | ** |
|  | STAT2 | 33.8 | 261.9 | 41.0 | 210.7 | 16.3 | 52.8 | ** |
|  | AIM2 | 0 | 10.0 | 0 | 6.6 | 0 | 2.2 | * |
|  | CASP1 | 0 | 56.0 | 0.5 | 21.8 | 0.6 | 3.8 | ** |
| Midkine, Chemokines, & Cytokines | MDK | 599.6 | 2058.9 | 543.2 | 990.9 | 330.5 | 332.2 | * |
|  | LGALS9 | 0.07 | 112.23 | 0.08 | 59.30 | 0.21 | 11.36 | ** |
|  | CXCL11 | 0.1 | 231.5 | 0 | 54.2 | 1.0 | 47.3 | ** |
|  | CXCL10 | 0.1 | 137.2 | 0 | 48.7 | 0.3 | 29.9 | ** |
|  | IFNL1 | 0 | 131.1 | 0 | 33.8 | 0.2 | 26.7 | ** |
|  | IFNB1 | 0 | 113.1 | 0 | 21.0 | 0 | 16.3 | ** |
|  | IFNL2 | 0 | 67.8 | 0.1 | 22.1 | 0.1 | 7.0 | * |
|  | IFNL3 | 0 | 41.2 | 0 | 16.6 | 0.2 | 5.3 | * |
|  | IFNA7 | 0 | 3.7 | 0 | 0.4 | 0 | 0 | * |
|  | IFNA10 | 0 | 1.5 | 0 | 0.1 | 0 | 0 | * |
|  | IFNA16 | 0 | 0.8 | 0 | 0.2 | 0 | 0 | * |
| Selected CSGs | BST2 | 3.1 | 1145.8 | 0.6 | 652.8 | 0.1 | 136.3 | ** |
|  | PSMB9 | 5.8 | 191.7 | 7.8 | 196.3 | 4.8 | 39.6 | ** |
|  | IFIT2 | 23.1 | 3070.6 | 44.4 | 1916.0 | 26.4 | 347.8 | ** |
|  | TRIM22 | 0.77 | 218.36 | 0.10 | 105.61 | 0.06 | 17.53 | ** |
|  | RSAD2 | 2.9 | 1418.0 | 0.9 | 768.7 | 1.3 | 185.6 | ** |
|  | LGALS3BP | 84.7 | 1313.9 | 163.5 | 1512.9 | 94.9 | 336.8 | ** |
|  | SERPINE1 | 90.4 | 789.8 | 285.9 | 1854.8 | 164.0 | 134.0 | ** |
|  | CD68 | 50.6 | 727.2 | 88.5 | 439.1 | 55.5 | 109.4 | ** |
|  | WARS1 | 98.8 | 704.1 | 119.5 | 256.3 | 99.1 | 129.1 | ** |
|  | SAMD9 | 12.1 | 665.7 | 14.5 | 347.7 | 11.8 | 149.2 | ** |

**Gene expression levels significantly lower in electroporated PC-3 cells versus cells Lipofected in SFM and SCM

*Gene expression levels significantly lower in electroporated PC-3 cells only compared to cells Lipofected in SFM (i.e., not significantly lower than PC-3 cells Lipofected in SCM)
