## Supplementary Information for "Transcriptomic Analysis of the Innate Immune Response to *in vitro* Transfection of Plasmid DNA"

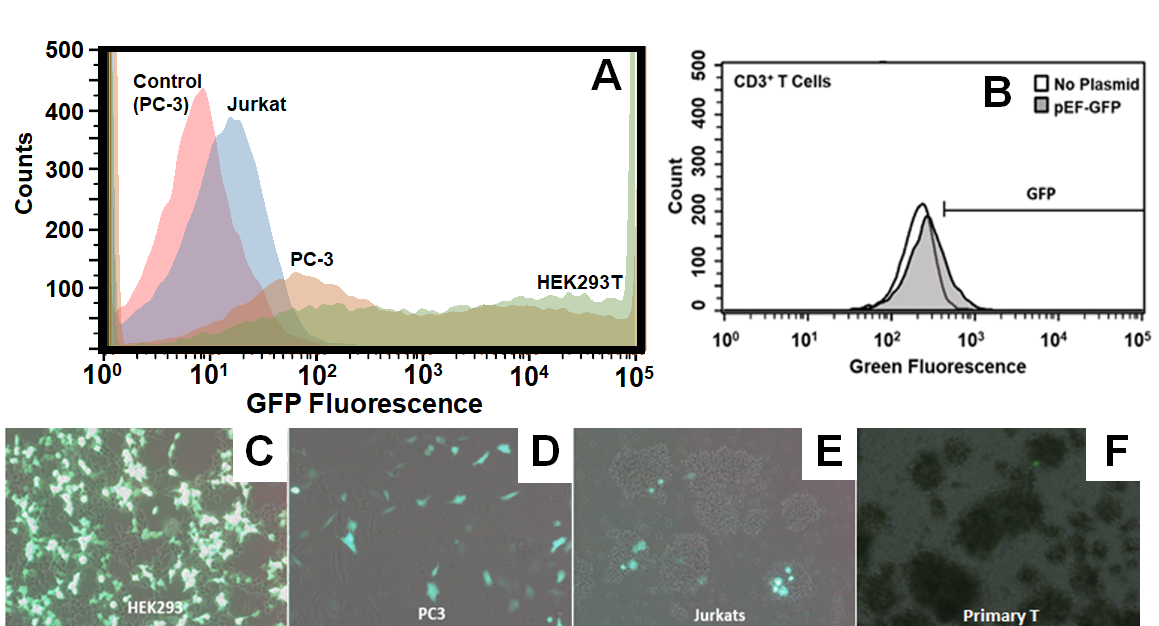

**Figure S1** – Representative histograms for transfected Jurkat, PC-3, HEK-293T cells (A) and primary T cells (B) compared to untransfected control cells. (C-F) Fluorescent microscopy images of GFP expression in each cell line 24 hours after transfection.

*#Load the counts data as a matrix. The counts file should be saved as a tab-delimited .txt file.*

cts <- as.matrix(read.csv('cts.txt',sep="\t",row.names="gene_id"))

*#Load the coldata matrix, which defines which samples will be grouped and compared. The file should be saved as a .csv. Columns = sample name, condition, and type. Sample names/rows should be the same as column names in the counts file.*

coldata <- read.csv("coldata.csv", row.names=1)

coldata <- coldata[,c("condition","type")]

coldata$condition <- factor(coldata$condition)

coldata$type <- factor(coldata$type)

*# Call the DESeq2 routine for analysis and build a dataset (dds)*

library("DESeq2")

dds <- DESeqDataSetFromMatrix(countData = cts, colData = coldata, design = ~ condition)

dds

library("DESeq2")

*#Run the analysis*

dds <- DESeq(dds)

res <- results(dds, independentFiltering = FALSE)

res

*#Order results by adjusted p-value*

resOrdered <- res[order(res$pvalue),]

*#Export results as a DEG.csv file*

write.csv(as.data.frame(resOrdered), file="DEG.csv")

Figure S2 – Script used to analyze mRNA-sequencing counts with DESeq2 in R Studio.

**Table S1** – Sequences of the qPCR primers used in Figure 4

| Target | Forward Primer | Reverse Primer |
| --- | --- | --- |
| IFNB1 | GAAGGAGGACGCCGCATTGA | TGCTCATGAGTTTTCCCCTGGT |
| IFNL1 | GGTGACTTTGGTGCTAGGCT | TGAGTGACTCTTCCAAGGCG |
| IFNL2 | GGGTGACAGCCTCAGAGTGTT | ACTCTTCTAAGGCATCTTTGGCCC |
| IFNL3 | TGAAACTAGACATGACCGGGGAC | CGGCACTTGCAGTCCTTCAG |
| CXCL10 | CCACGTGTTGAGATCATTGCT | TGCATCGATTTTGCTCCCCT |
| CXCL11 | AGTCCTGGAAAAGGGCATCTG | TTTGGTCCTTTCACCCACCT |
| CASP1 | AATACTGTCAAATTCTTCATTGCAGATAAT | AAGTCGGCAGAGATTTATCCAATAA |
| GAPDH | CAATGACCCCTTCATTGACC | GACAAGCTTCCCGTTCTCAG |

**Table S2.** Expression levels (TPM) of chemokines and cytokines in transfected (Trans) and untransfected (Ctrl) cells.

|  | HEK-293T | | PC-3 | | Jurkat | | Primary T | |
| --- | --- | --- | --- | --- | --- | --- | --- | --- |
|  | Ctrl | Trans | Ctrl | Trans | Ctrl | Trans | Ctrl | Trans |
| CXCL1 | 0.1 | 0.2 | 54.4 | 214.2 | 0 | 0 | 0.3 | 0.2 |
| CXCL2 | 0.6 | 0.7 | 2.1 | 40.0 | 0 | 0 | 0.1 | 0.2 |
| CXCL3 | 0.4 | 0.9 | 3.2 | 47.9 | 2.3 | 1.7 | 0.2 | 0.1 |
| CXCL5 | 0 | 0.1 | 3.1 | 6.6 | 0 | 0 | 0 | 0 |
| CXCL6 | 0 | 0.1 | 24.3 | 65.4 | 0 | 0 | 0.1 | 0 |
| CXCL8 | 0.3 | 0.2 | 56.2 | 608.7 | 0 | 0 | 125.2 | 379.9 |
| CXCL9 | 0 | 0.0 | 0 | 2.1 | 0 | 0 | 0.9 | 1.6 |
| CXCL10 | 0 | 0.1 | 0.1 | 137.2 | 0 | 0 | 21.0 | 24.5 |
| CXCL11 | 0 | 0 | 0.1 | 231.5 | 0 | 0 | 2.9 | 4.0 |
| CXCL13 | 0 | 0 | 0 | 0.0 | 0 | 0 | 33.1 | 33.6 |
| IFNα7 | 0 | 0.1 | 0 | 3.7 | 0 | 0 | 0 | 0 |
| IFNα10 | 0 | 0 | 0 | 1.5 | 0 | 0 | 0 | 0 |
| IFNα13 | 0 | 0 | 0.1 | 1.3 | 0 | 0 | 0 | 0 |
| IFNβ1 | 0 | 0 | 0 | 113.1 | 0 | 0 | 0.1 | 0.1 |
| IFNλ1 | 0 | 0 | 0 | 131.1 | 0 | 0 | 0.8 | 0.5 |
| IFNλ2 | 0 | 0.1 | 0 | 67.8 | 0 | 0 | 0 | 0 |
| IFNλ3 | 0 | 0 | 0 | 41.2 | 0 | 0 | 0 | 0 |
| IL1β | 0 | 0 | 24.2 | 56.9 | 0 | 0 | 0.6 | 0.3 |
| IL6 | 0 | 0 | 1.2 | 130.0 | 0 | 0 | 0.4 | 0.5 |
| IL18 | 0.1 | 0 | 91.5 | 143.7 | 0 | 0 | 0.5 | 0.3 |
| TNFα | 0 | 0 | 0 | 0.7 | 0 | 0 | 547.8 | 751.8 |

**Table S3.** Expression levels of select CSGs in transfected (Trans) and untransfected (Ctrl) cells.

| Gene  Symbol | HEK-239T | | PC-3 | | Jurkat | | Primary T | |
| --- | --- | --- | --- | --- | --- | --- | --- | --- |
|  | Ctrl | Trans | Ctrl | Trans | Ctrl | Trans | Ctrl | Trans |
| BST2 | 0.7 | 1.0 | 3.1 | 1145.8 | 97.3 | 103.7 | 261.8 | 251.8 |
| IFI16 | 0.0 | 0.0 | 25.7 | 945.5 | 294.1 | 266.8 | 274.9 | 215.1 |
| PSMB8 | 0.1 | 0.2 | 35.4 | 361.1 | 232.0 | 247.7 | 353.7 | 305.2 |
| PSMB9 | 0.1 | 0.3 | 5.8 | 191.7 | 116.7 | 124.9 | 182.4 | 171.0 |
| IRF1 | 3.0 | 3.5 | 17.1 | 139.7 | 73.7 | 75.8 | 65.1 | 62.1 |
| ISG15 | 5.4 | 6.3 | 181.5 | 8355.9 | 161.8 | 166.1 | 61.3 | 63.1 |
| OASL | 0.0 | 0.0 | 9.6 | 2137.8 | 1.3 | 1.4 | 25.1 | 19.6 |
| OAS2 | 0.0 | 0.0 | 39.1 | 1596.4 | 19.2 | 18.0 | 78.6 | 72.7 |
| IL32 | 0.7 | 0.7 | 14.0 | 55.2 | 350.5 | 427.4 | 1613.7 | 1062.2 |
| OAS1 | 0.0 | 0.0 | 35.0 | 1490.6 | 2.8 | 2.7 | 27.8 | 20.3 |
| UBE2L6 | 3.0 | 3.3 | 45.5 | 1052.8 | 161.8 | 181.0 | 170.9 | 161.2 |
| OAS3 | 0.1 | 0.2 | 53.3 | 1129.6 | 17.3 | 17.6 | 40.2 | 32.2 |
| IL2RG | 0.0 | 0.0 | 0.5 | 4.0 | 396.6 | 414.4 | 824.0 | 624.7 |
| LAMP3 | 0.7 | 1.1 | 19.6 | 738.5 | 43.4 | 41.7 | 4.9 | 4.9 |
| SAMD9 | 0.3 | 0.3 | 12.1 | 665.7 | 19.7 | 17.2 | 21.6 | 12.9 |
| IFIH1 | 0.4 | 0.6 | 10.7 | 593.2 | 9.8 | 8.8 | 19.0 | 18.8 |
| PARP9 | 0.9 | 1.4 | 44.2 | 487.8 | 20.9 | 19.2 | 33.9 | 30.8 |
| IFI44 | 0.0 | 0.0 | 20.2 | 507.7 | 2.1 | 2.3 | 16.5 | 16.4 |
| SP110 | 0.2 | 0.3 | 16.2 | 386.7 | 14.7 | 13.2 | 62.3 | 65.6 |
| NMI | 1.6 | 1.7 | 35.0 | 245.2 | 152.9 | 133.8 | 73.1 | 64.0 |
| CD7 | 0.1 | 0.2 | 0.6 | 14.2 | 220.6 | 259.8 | 216.3 | 163.6 |
| MYD88 | 1.3 | 1.9 | 52.9 | 298.3 | 77.8 | 84.3 | 60.9 | 48.4 |
| ICAM1 | 0.0 | 0.7 | 40.5 | 320.4 | 1.5 | 1.5 | 85.7 | 96.8 |
| DTX3L | 0.8 | 1.1 | 33.6 | 329.6 | 30.9 | 28.7 | 31.9 | 26.8 |
| GBP1 | 0.1 | 0.2 | 2.4 | 298.5 | 24.5 | 20.1 | 59.3 | 55.9 |
| SHFL | 0.6 | 0.6 | 17.9 | 308.3 | 29.5 | 29.9 | 42.6 | 33.9 |
| SAMD9L | 0.0 | 0.0 | 5.3 | 354.5 | 2.8 | 2.2 | 17.7 | 12.1 |
| SELL | 0.0 | 0.1 | 0.3 | 4.5 | 271.4 | 246.0 | 342.3 | 110.0 |
| PARP14 | 0.0 | 0.0 | 31.7 | 306.9 | 20.6 | 18.5 | 42.6 | 34.7 |
| GSDMD | 0.0 | 0.0 | 72.8 | 245.4 | 59.3 | 62.2 | 45.1 | 37.9 |
| LGALS9 | 0.9 | 0.8 | 0.1 | 112.2 | 124.7 | 137.1 | 14.5 | 10.1 |
| ERAP1 | 2.4 | 3.5 | 36.6 | 130.4 | 60.2 | 55.9 | 71.7 | 63.4 |
| STING1 | 0.2 | 0.6 | 19.0 | 42.2 | 56.6 | 59.9 | 144.1 | 141.7 |
| TRIM14 | 3.9 | 6.5 | 25.6 | 118.7 | 80.3 | 81.6 | 44.5 | 32.4 |
| FYB1 | 0.0 | 0.0 | 0.3 | 9.8 | 177.8 | 153.3 | 87.5 | 51.5 |
| MVP | 0.4 | 0.6 | 24.3 | 67.8 | 8.6 | 9.7 | 137.1 | 124.3 |
| APOBEC3G | 0.1 | 0.2 | 28.5 | 85.2 | 6.4 | 6.5 | 132.4 | 105.8 |
| BIRC3 | 0.1 | 0.0 | 7.7 | 53.0 | 10.2 | 8.7 | 118.6 | 128.5 |
| ARHGAP15 | 0.0 | 0.0 | 0.0 | 1.4 | 117.0 | 105.5 | 102.2 | 72.1 |
| SP140L | 0.0 | 0.1 | 15.4 | 105.9 | 31.7 | 28.9 | 41.2 | 31.0 |
| XAF1 | 0.0 | 0.0 | 5.2 | 131.5 | 9.3 | 9.0 | 24.2 | 18.5 |
| SLFN5 | 0.6 | 0.8 | 19.3 | 130.9 | 19.1 | 18.0 | 31.1 | 9.5 |
| APOL6 | 0.0 | 0.0 | 9.9 | 100.4 | 16.7 | 14.5 | 38.4 | 36.9 |
| PHF11 | 0.5 | 0.3 | 13.2 | 89.1 | 36.9 | 34.2 | 24.1 | 23.7 |
| TRIM56 | 0.9 | 1.1 | 10.4 | 54.1 | 62.5 | 63.8 | 24.9 | 19.5 |
| PARP12 | 0.0 | 0.0 | 6.9 | 98.6 | 21.2 | 22.8 | 14.0 | 11.0 |
| IL7R | 0.0 | 0.0 | 5.8 | 73.8 | 11.1 | 9.0 | 68.4 | 27.2 |
| GIMAP2 | 0.0 | 0.0 | 0.0 | 4.7 | 88.8 | 82.6 | 34.0 | 22.6 |
| CD96 | 0.1 | 0.1 | 0.5 | 2.3 | 23.9 | 21.5 | 135.0 | 84.9 |
| TNFRSF14 | 0.0 | 0.1 | 9.4 | 41.4 | 10.4 | 12.3 | 40.7 | 41.7 |
| APOL3 | 0.0 | 0.0 | 0.8 | 42.8 | 7.8 | 6.8 | 46.0 | 27.2 |
| CLEC2B | 0.0 | 0.0 | 7.3 | 25.0 | 6.5 | 5.6 | 27.0 | 21.2 |
| TMEM229B | 0.5 | 0.6 | 0.5 | 15.6 | 15.6 | 14.5 | 2.1 | 1.9 |
| ATP10A | 0.0 | 0.0 | 0.6 | 7.3 | 3.2 | 3.5 | 23.3 | 19.5 |

**Table S4.** Expression levels (TPM) of HSPGs in primary T cells, Jurkat T cells, PC-3 cells, and HEK293-T cells.

|  | HEK-293T Cells | | PC-3 Cells | | Jurkat T Cells | | Primary T Cells | |
| --- | --- | --- | --- | --- | --- | --- | --- | --- |
| Gene | Control | Trans. | Control | Trans. | Control | Trans. | Control | Trans. |
| HSPG2 | 2.3 | 2.4 | 41.9 | 34.2 | 0.05 | 0.07 | 0.3 | 0.1 |
| SDC1 | 3.9 | 6.7 | 173.1 | 71.9 | 0 | 0 | 0 | 0 |
| SDC2 | 32.4 | 36.0 | 30.3 | 15.1 | 0 | 0 | 0.1 | 0.1 |
| SDC3 | 15.3 | 22.4 | 12.9 | 9.1 | 8.1 | 9.3 | 0 | 0.1 |
| SDC4 | 32.5 | 26.0 | 70.6 | 103.4 | 0 | 0 | 194.1 | 203.7 |
